# The Gompertz curve for estimating growth rates of Protein Data Bank and protein folds

**DOI:** 10.64898/2026.06.24.732253

**Authors:** Kochi Sato, Kentaro Tomii

## Abstract

The Protein Data Bank (PDB) is an ever-growing, open-access repository of structural data of biological molecules. This international database has been instrumental in the development of artificial intelligence and deep learning models for protein structure prediction and design. The PDB growth is a crucially important factor influencing further development of these models. Therefore, after analyzing the growth trend in PDB depositions since the archive’s launch, we found that it is well fitted by the Gompertz function, a growth curve used across various disciplines. Furthermore, we observed that the function captures the “discovery of novel folds”, i.e., the cumulative number of distinct folds among protein domains that constitute most of the PDB. Consequently, based on the fitting results, we estimated the likely numbers of PDB entries and protein folds. These findings provide insights into deceleration of growth in recent years and enable us to assess anticipated trends.

## Introduction

As a global core biodata resource, the Protein Data Bank (PDB) is indispensable for providing free and public access to three-dimensional (3D) structural data on biological molecules, offering strong and continuous support to biological and biomedical research^1^. The PDB serves as the repository of experimentally determined 3D atomic coordinates of proteins/peptides and nucleic acids with and without ligands, as well as their assemblies. Those coordinates are determined by X-ray crystallography, nuclear magnetic resonance (NMR) spectroscopy, cryogenic electron microscopy (cryo-EM), and other methods. Determined and deposited structures range from monomeric proteins such as myoglobin to supramolecular complexes of proteins and nucleic acids such as ribosomes and nucleosomes. The PDB has driven innovations in biological and biomedical research, enabling new methods of scientific discovery such as AlphaFold^2^. Going beyond those “experimentally-determined, rigorously validated, and expertly-biocurated” structures, Computed Structure Models (CSMs) have been incorporated from external repositories, including AlphaFoldDB^3,4^. Since its inception in 1971 at Brookhaven National Laboratory^5^, when only seven structures were available, the number of PDB entries has grown remarkably. In fact, as of March 2026, the number of PDB depositions has reached over 250,000 experimentally determined entries.

While the PDB has been growing rapidly overall, the growth rate has been non-monotonic and has notably slowed in recent years^6^. Accordingly, several growth models have been proposed to analyze and interpret the changes of the growth rate. At a very early stage, Dickerson, in his renowned 1978 letter^7^, described the growth as exponential, represented as (1/0.19)exp[0.19(*t*/yr − 1960)], where *t* denotes the calendar year. Although this model captured the archive size well until the 1990s, systematic deviations emerged thereafter. To accommodate the changing rate, Levitt^6^ proposed a compound model with a power law: 21.9(*t*/yr − 1987)^2.17^exp[0.062(*t*/yr − 1987)]. According to Abad-Zapatero’s reanalysis based on the crystallographic data through 2011, the exponential factor was estimated as 0.252. However, as described there, “a definitive downward trend after 2001” was also apparent despite enormous efforts and investments in structural determinations^8^.

Here, we propose a Gompertz function as an alternative growth curve of PDB entries, and also present the perspectives and prospects based on the fitting results over a half century from the first deposition. In addition, we demonstrate the effectiveness of the fitting approach for assessing the growth of novel protein folds, whose upper limit is one of the most important questions in structural biology. The Gompertz function, first introduced by Gompertz to model the human mortality rate^9^, is defined as follows:

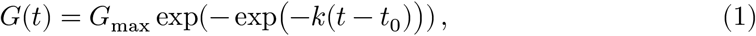

where parameters *G*_max_, *k*, and *t*_0_ represent the maximum value, the growth rate constant, and the inflection point, respectively. This function describes the model with the relative growth rate declining exponentially, which often captures the growth curve for groups with a resource limitation such as biological growth and epidemic spread^10,11^. The curve shows asymmetric sigmoidal behavior: a super-exponential increase in the initial phase, exponential growth near the inflection point, and convergence toward saturation at the end. These results shed light on the interpretation of the PDB growth and give us some clues to estimate future trends and developments.

## Results

### Results of Global Fitting

The number of PDB entries has not continued the exponential growth observed in its early stage, and in recent years both the Dickerson and Levitt curves have tended to overestimate the PDB growth curve. In fact, although the number of depositions to the PDB reached 250,000 in 2026, the Dickerson and Levitt curves overestimate the archive size by factors of approximately 4.8 and 2.4, respectively.

Thus, we applied the Gompertz function to the growth curve of PDB entries per year from 1971, when the first entry was submitted, to 2025. In contrast to the models described above, the Gompertz curve provides a good fit to the observed data, with a low root mean square error (RMSE), closely approximating the growth curve (Fig. 1A). Unlike the conventional monotonic growth models, it yields an S-shaped trajectory. The fitting results show that the inflection point (*t*_0_) lies around the present time, indicating a decline in the growth rate over the coming decades (Table 1). The upper asymptote (*G*_max_) was estimated to be 8.4×10^5^, corresponding to approximately 3–4 times the current number of deposited entries.

**Table 1.** Parameters of the Gompertz function.

| | $G_{\max}$ | $k / \text{yr}^{-1}$ | $t_0 / \text{yr}$ |
| --- | --- | --- | --- |
| PDB | $(8.41 \pm 0.45) \times 10^5$ | $(4.74 \pm 0.13) \times 10^{-2}$ | $2029.6 \pm 1.0$ |
| X-ray | $(4.34 \pm 0.04) \times 10^5$ | $(6.26 \pm 0.05) \times 10^{-2}$ | $2020.9 \pm 0.2$ |
| NMR | $(1.63 \pm 0.02) \times 10^4$ | $(1.15 \pm 0.03) \times 10^{-1}$ | $2006.1 \pm 0.1$ |
| Cryo-EM | $(9.9^{+4.8}_{-3.3}) \times 10^5$ | $(7.35 \pm 0.71) \times 10^{-2}$ | $2041.8 \pm 3.2$ |
| ECOD X-group | $(3.61 \pm 0.09) \times 10^3$ | $(8.47 \pm 0.25) \times 10^{-2}$ | $2007.0 \pm 0.4$ |
| SCOPe fold | $(4.695 \pm 0.001) \times 10^3$ | $(6.54 \pm 0.88) \times 10^{-2}$ | $2011.1 \pm 3.8$ |
| mpstruc (unique) | $(2.7 \pm 1.2) \times 10^4$ | $(3.80 \pm 0.51) \times 10^{-2}$ | $2049.8 \pm 7.9$ |

**Figure 1.**
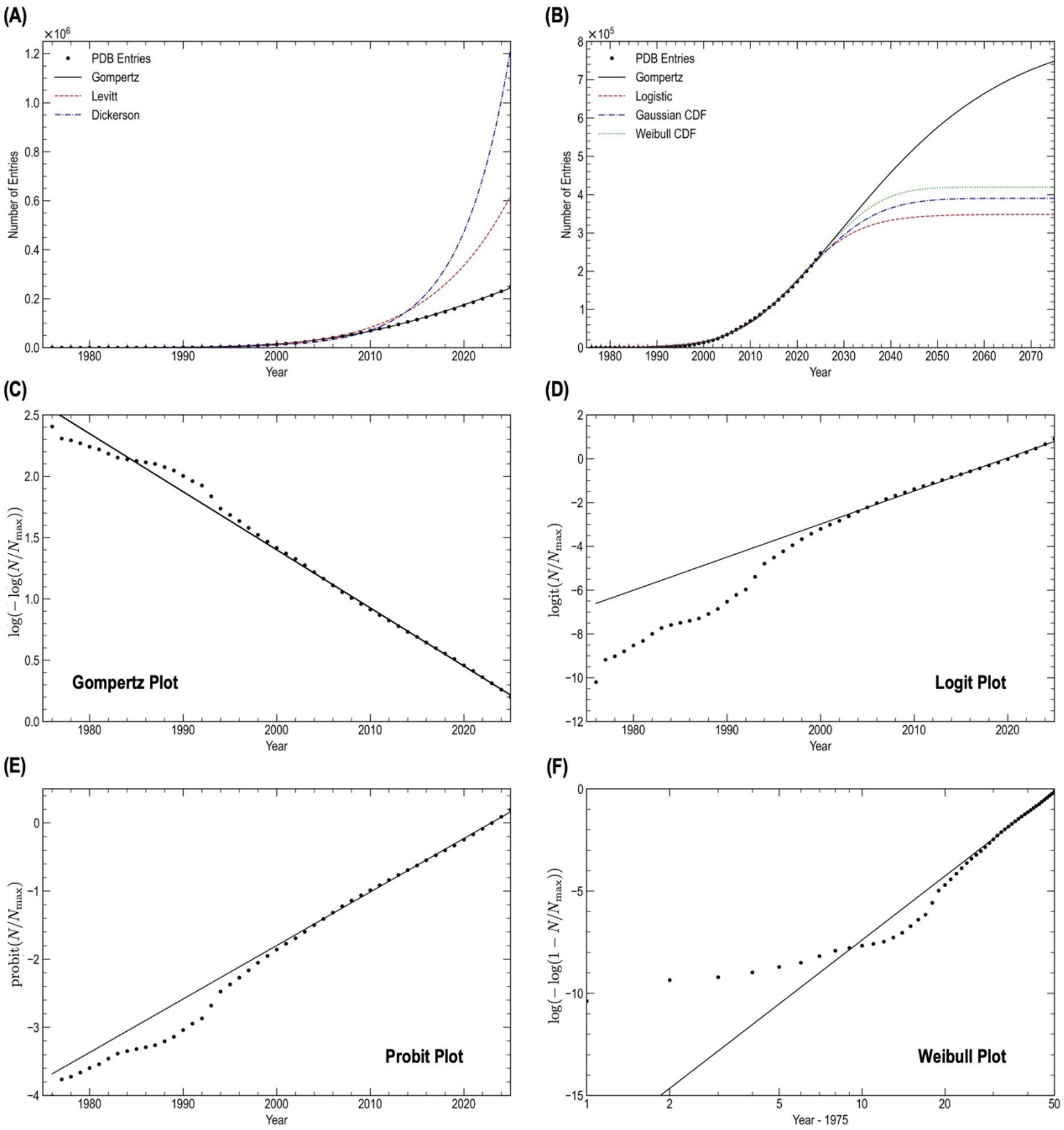
Growth curve of the PDB archive and comparison with representative growth models. (A) Growth curve of the total PDB entries (black dots) fitted with the Gompertz function (black line), compared with the conventional models proposed by Levitt (red dashed line) and Dickerson (blue dash–dotted line). (B) Comparison of the Gompertz fit (black) with other sigmoidal functions, including logistic function (red), Gaussian CDF (blue), and Weibull CDF (green). (C–F) Linear plots of the PDB growth curve, shown as (C) Gompertz plot, (D) logit plot, (E) probit plot, and (F) Weibull plot.

### Comparison with Other Functions

To assess whether the observed curve follows a Gompertz-type growth pattern, we compared the fitting performance with other S-shaped functions that are widely used to describe growth processes, such as the logistic function, the Gaussian cumulative distribution function (Gaussian CDF), and the Weibull cumulative distribution function (Weibull CDF) (see the *Methods*). When fitting globally to the entire growth curve, clear differences were observed in their behavior including fitting performance, inflection points, and upper asymptotes (Fig. 1B, Table 2, Supplementary Tables S1–S3). The logistic curve fails to reproduce a rapid increase in the initial region, and thus overestimates the data before 2000 and underestimates the values around 2010, resulting in a poor fit to the growth curve (Supplementary Fig. S1). The Gaussian CDF shows smaller deviations, but still overestimates the early-stage data and tends to underestimate the most recent observations since 2020. The Weibull CDF appears to capture the overall shape of the growth curve within the fitted range, as does the Gompertz function.

**Table 2.** Summary of the fitting results.

|  | RMSE | AIC | Estimated Limit |
| --- | --- | --- | --- |
| Gompertz | $1.34 \times 10^3$ | $1.45 \times 10^3$ | $(8.41 \pm 0.45) \times 10^5$ |
| Logistic | $3.21 \times 10^3$ | $1.62 \times 10^3$ | $(3.48 \pm 0.14) \times 10^5$ |
| Gaussian CDF | $1.94 \times 10^3$ | $1.52 \times 10^3$ | $(4.29 \pm 0.19) \times 10^5$ |
| Weibull CDF | $1.73 \times 10^3$ | $1.50 \times 10^3$ | $(4.19 \pm 0.24) \times 10^5$ |

Focusing on future projections, all models other than the Gompertz curve predict a rapid decline in growth rate in the near term, leading to an unrealistically early plateau within approximately 15 years. Moreover, their upper limits were about half of the estimates by the Gompertz curve, corresponding to only 1.7 times the current number of deposited entries. Consistent with these observations, the fitting metrics indicated that the Gompertz curve provides the best fit among the tested models, exhibiting lower RMSE and Akaike information criterion (AIC) than the other S-shaped functions (Table 2).

### Validation by Linearization

To validate the growth curve fittings, we constructed several linearized plots (Figs. 1C–F), including the Gompertz plot, logit plot, probit plot, and Weibull plot (see the *Methods*; Supplementary Notes S1–S2). In the Gompertz plot with the estimated *G*_max_, the PDB deposition data fall approximately along a straight line, supporting the validity of the Gompertz growth model. The probit plot also appears roughly linear over the entire range. In contrast, the logit plot shows clear deviations from linearity over the first two decades, indicating that the logistic function does not adequately describe the PDB growth. The Weibull plot shows substantial deviations from linearity across most of the range. These linear transformations collectively support the Gompertz model, rather than the logistic function and the Weibull CDF.

We next examine the systematic deviation patterns from linearity across the linearized plots. In the Gompertz plot, the data fluctuate around the fitted line in the early region and converge at later times, whereas in the logit and probit plots they lie below the fitted line over the first half and approach it only later. The Weibull plot deviates substantially across the entire range. Given that these transformations are monotonic, the consistent downward deviation in the logit and probit plots indicates that the initial values are lower than those predicted by the logistic and Gaussian CDF models, followed by a delayed but steeper rise. Together with the agreement in the Gompertz plot, the lack of overshoot and subsequent convergence further demonstrates that the growth is not symmetric but biased toward later times.

### Interpretation of Early Growth Behavior

As described above, the PDB growth exhibits a very sharp initial rise. In the Weibull model, the early-time behavior follows a finite power law, where the order is given by the shape parameter *m*, corresponding to the slope of the Weibull plot and estimated to be approximately 4.6 globally and close to 1 in the initial regime. By contrast, the logistic function exhibits exponential growth, whereas the Gaussian CDF has its onset governed by a Gaussian tail. The Gompertz model, however, displays a super-exponential rise in the initial regime. These comparisons indicate that the observed onset of PDB growth is more abrupt than can be accounted for by polynomial (Weibull), exponential (logistic), or Gaussian-tail (Gaussian CDF) behavior, and is instead consistent with a super-exponential form as realized by the Gompertz model.

This raises the question of why early observations, such as those by Levitt and Dickerson, appeared consistent with exponential growth. Notably, neither the Gaussian CDF nor the Weibull model exhibits an exponential-like regime. In contrast, in the Gompertz model, when the absolute count is vanishingly small relative to the upper limit, the super-exponential growth becomes effectively indistinguishable from simple exponential growth. In this regime (1 < *G* ≪ *G*_max_), the relative growth rate reduces to an approximately constant value:

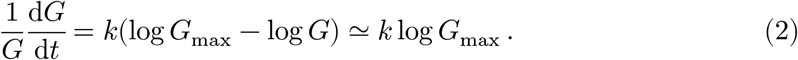

This initial growth behavior accounts for the exponential growth highlighted by Dickerson and Levitt, furthermore, the estimated rate agrees in order of magnitude with Dickerson’s rate 0.19 (≃ *k* log *G*_max_) and Levitt’s rate 0.062 (≃ *k*).

### Gompertz Fitting by Experimental Methods

To further investigate the contributions of the structure determination methods to the growth curve, Gompertz fitting was applied to each method’s curve – X-ray crystallography, NMR, and cryo-EM – which are mainly employed for structural analysis. As well as the total deposited entries, all these method-specific curves were closely approximated by Gompertz functions (Figs. 2A–C, Table 1). The growth curves of the three methods exhibit markedly different stages at present, with X-ray crystallography near an inflection point, NMR showing saturation, and cryo-EM still in an early phase. Although the maximum number of PDB entries determined by cryo-EM was estimated to be relatively high with substantial uncertainty, the sum of the estimated upper limits for all methods was on the order of 10^6^, almost consistent with the estimated maximum of the total growth curve.

**Figure 2.**
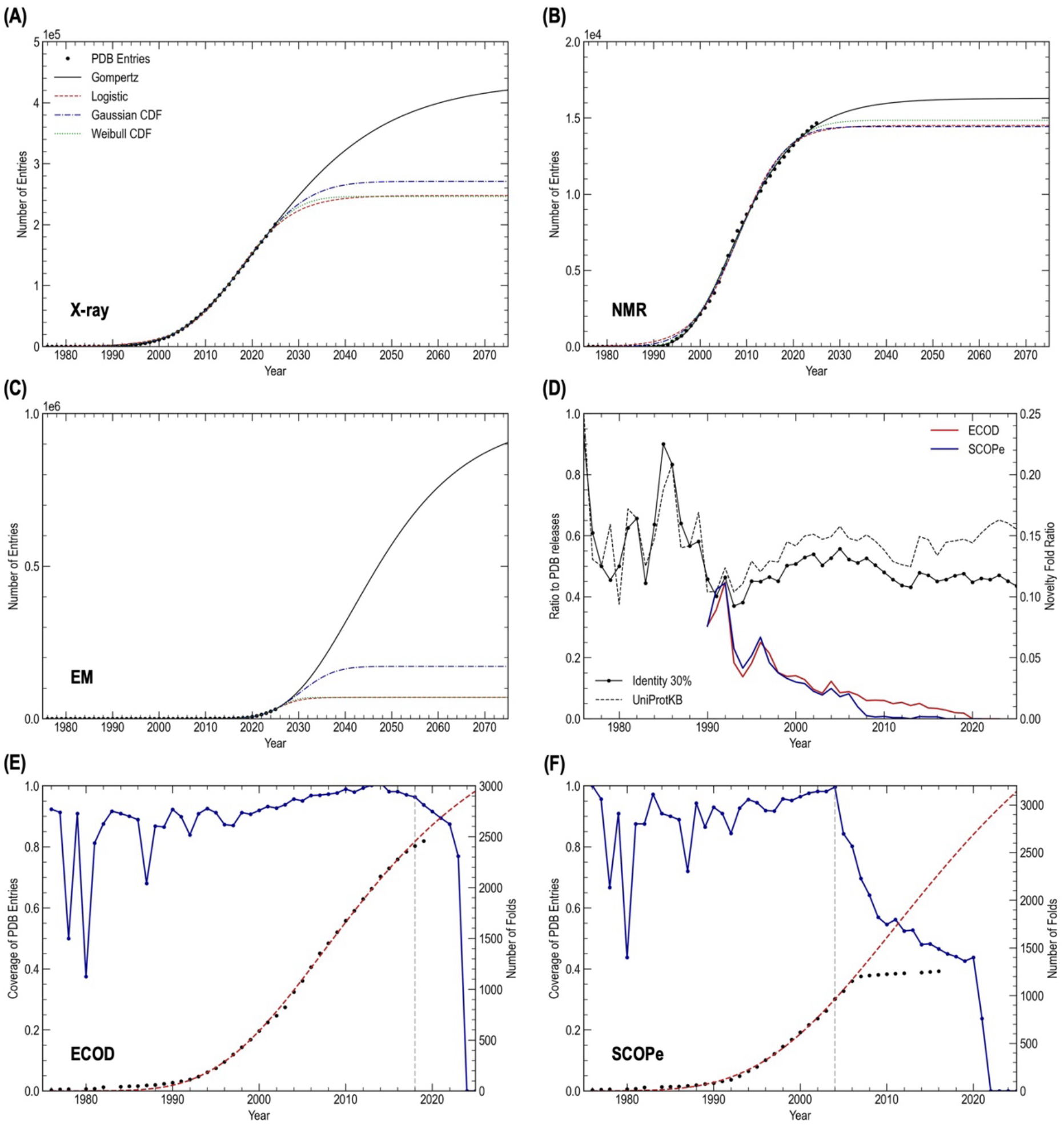
Growth curves of the PDB archive for structure determination methods and structural categories. (A–C) Comparisons of fitted curves using Gompertz (black line), logistic (red dashed line), Gaussian CDF (blue dash–dotted line), and Weibull CDF (green dotted line) for (A) X-ray structures, (B) NMR structures, and (C) EM structures. (D) Novelty in the PDB is shown as the number of releases 30% identity clusters (black dots) and unique UniProtKB entries (black dashed line), overlaid with the fold-level novelty ratios from ECOD (red line) and SCOPe (blue line). (E–F) Growth curves of (E) ECOD X-groups and (F) SCOPe folds (black dots) are shown with their coverage relative to the PDB archive (blue dots/line). Gompertz curves fitted to the fold growth are shown as red dashed lines, with the fitting cutoff shown by a grey vertical line.

In comparison, all the other models than the Gompertz exhibit systematic inconsistencies when applied to the method-specific growth curves. For X-ray crystallography, these models predict saturation within only a decade (Fig. 2A). In the case of NMR, all non-Gompertz models yield estimated upper limits that already fall below the current count (Fig. 2B). Similarly, for cryo-EM, these models again predict near-term saturation, with estimated upper limits of no more than 70,000 for the logistic and Weibull models, approximately twice the current level, and only 170,000 for the Gaussian model (Fig. 2C). In all cases, prominently for NMR, the fitted curves tended to underestimate the growth at later times while overestimating the initial increase. This behavior is consistent with that observed in the global fitting described above, reflecting a rapid initial rise and an asymmetric growth curve with a longer tail toward later times. In other words, fitting to the steep initial rise leads to premature saturation, whereas fitting to the later growth results in a broadened initial profile.

### Novelty Ratios and Folds

The number of PDB entries is increasing as described above, however some novelty ratios show a decreasing trend. While unique cluster ratios to the annual released PDB structures, such as unique protein sequences at identity 30% and unique UniProtKB entries with known 3D structures, have remained nearly constant at 10–20%, the novelty ratios of fold categories have been declining for both ECOD (Evolutionary Classification of protein Domains) X-groups (possible homology)^12^ and SCOPe (Structural Classification of Proteins – extended) fold groups^13^, derived from structural classifications of protein domains (Fig. 2D). As in the case of the PDB entries, growth curves of those categories were also approximated by Gompertz curves (Figs. 2E and 2F, Table 1). The discovery of novel folds seems to be converging in 2020s, and the maximum numbers were estimated to be approximately 3,600 in ECOD and 4,700 in SCOPe.

## Discussion

The PDB growth is essential for progress in biological and biomedical research, especially for the development of methods to predict and design biomolecular structures. Estimating anticipated data growth is a key to the further development of artificial intelligence and deep learning models that rely heavily on large and high-quality datasets. Looking ahead, the fitting results suggest that the number of entries will double in the next decade, mainly driven by the contributions of the structure determination by cryo-EM. However, the overall growth rate of the PDB data in terms of the number of entries is expected to slow, consistent with a field-wide transition from single-protein determinations to more complex targets such as supramolecular complexes and multicomponent assemblies. This consistency suggests that the growth curve reflects the world’s capacity to determine structures of biological macromolecules because the Gompertz function captures growth under resource limitations. Therefore, the fitting results appear to provide a natural framework for modeling archive maturation in structural biology, a field constrained by both human and technological resources. In other words, we speculate that the deposition rate is approaching the “maximum” rate at which the structural biology community can produce data for the PDB.

Although the estimated asymptotic limit might appear small compared with the possible number of proteins (whose structures remain to be determined), it is noteworthy that the complexity and/or number of molecules per PDB entry has been increasing over time.^1^ As the model does not account for the complexity or size of the contents, the upper asymptote does not necessarily represent the absolute maximum for the number of proteins. Therefore, we will have determined structures for considerably more molecules than 10^6^ in the future. This phenomenon may share some resemblance with the diffusion of technological innovations such as mobile telephony, where the devices involved underwent a similar evolution, becoming more sophisticated over time. These points suggest that the number of entries is an adequate metric for monitoring deposition rates, regardless of their complexity or size.

In addition, as described in *Gompertz Fitting by Experimental Methods*, the fitting results appear to serve as an indicator of the maturity of methodological and/or conceptual trends, as exemplified by stage differences across the three determination methods. The maturation observed in NMR may also be observed for X-ray and cryo-EM in the future. As often observed in computational technology maturation^14^ and trend life cycles in scientific research^15^, Gompertz curves might also be powerful for assessing the maturity of method-specific or subfield-specific trends. More broadly, the same approach may be applicable to characterizing the sizes of other scientific archives and to forecasting the maturity of science data more generally.

As an illustration, we considered a pseudo growth curve for a subfield focused on membrane proteins. mpstruc is a database of unique three-dimensional structures of membrane proteins, curated from PDB entries that have been published in peer reviewed journals^16^. The number of entries in mpstruc has steadily increased over time, with more than 10,000 deposited structures and over 2,000 unique entries currently available. The growth appears to slow, but it may reflect time lags associated with manual curation and publication. In fact, in the Gompertz plot, the number of unique structures follows an approximately linear relation, suggesting that the growth is well described by a Gompertz-type curve (Supplementary Fig. S2). A Gompertz fit to the growth curve up to 2025 yields an inflection point around 2050 and an estimated upper limit of approximately 30,000 unique clusters, indicating that the structure determination of membrane proteins is still in an early stage of growth (Table 1).

Apparently, the Gompertz curve provides an appropriate description not only of PDB growth but also of the growth of fold categories in hierarchical classification databases such as ECOD and SCOPe, where annual data are available, similarly to other discovery processes such as software fault detection^17^. This is consistent with Levitt’s assertion that the novel folds discovery scales with the growth of PDB archive.^6^ More specifically, to model the relation between the number of discovered folds and the number of explored protein families, a stretched exponential form has been introduced for the fold probability distribution.^18^ For distributions with such rapidly decaying tails, considering the exploration process through structure determination naturally leads to Gompertz-type growth. Introducing the rarity *X* = − log λ for a fold with probability λ, one can show that, for a broad class of distributions in which the tail probability Pr(*X* > *x*) decays exponentially or faster, the maximum rarity defining the coverage frontier converges weakly to a Gumbel extreme value distribution (Supplementary Note S3). Accordingly, if the rarity increases with time, corresponding to exponential expansion of the explored domain, then the resulting fold coverage is expected to follow a Gumbel CDF, i.e., a Gompertz curve.

The inferred upper limit for the total number of protein folds, approximately 3,500– 5,000, exceeds Levitt’s estimate and lies within the broad range of previously reported values (455–10,000)^19^. It is also consistent with the estimate of about 4,000 folds obtained from an earlier study that assumed a stretched exponential distribution of fold probabilities and modeled the fold discovery rate based on early SCOP data^18^. In contrast, our estimate is derived from later observations under weak assumptions that do not specify the detailed form of the fold distribution but which instead assume exponential expansion of the explored domain. The consistency between these estimates, despite the different assumptions and data regimes, is noteworthy, though the earlier study further suggested that only about 2,000 folds are likely to occur in nature, with many others being unlikely. By contrast, 6,433 high-symmetry and 7,427 low-symmetry novel fold candidate sets were recently identified among structure models predicted by AlphaFold at the Encyclopedia of Domains (TED)^20^. The inferred values appear to be lower than these numbers. However, a recent comparative analysis of TED and Domain Parser for AlphaFold Models (DPAM) domains suggests that the number of novel folds may be lower, as a substantial proportion of low-symmetry candidates can be assigned to known ECOD categories.^21^

In summary, our findings revealed that the Gompertz functions well approximate the growth curves of the PDB archive and protein folds and provide maturity metrics with predictive utility. Since this growth model is a rough representation based on past trends, some room remains for more detailed and sophisticated refinement. For example, rigorously, “it will be true that Gompertz curves will often add to give something very close to a Gompertz curve”. More importantly, technological innovations or topic shifts, such as a recent expansion of structure determinations using cryo-EM, may give rise to new growth regimes, adding uncertainty to future projections, as mentioned in *Gompertz Fitting by Experimental Methods*.

## Methods

### PDB and protein folds statistics

We obtained annual Protein Data Bank (PDB) depositions, method-specific PDB depositions, 30% sequence identity clusters, and unique UniProtKB entries from the PDB statistics available as of March 2026^1^. These annual counts were used to calculate PDB growth curves and to estimate the novelty ratio as a scale factor.

To deal with protein fold growth curves, we analyzed statistics of ECOD^12^ and SCOPe^13^, which are among the protein fold databases actively updated in recent years, and retrieved all deposited entries from the latest versions available as of March 2026 (ECOD ver. ver. 291* and SCOPe ver. 2.08). Specifically, we estimated the fold growth curves by counting the first PDB deposition dates in each fold category (i.e. x-groups in ECOD and fold groups in SCOPe), and noted that the novelty counts were treated as deposition counts in PDB rather than in fold databases to avoid potential artifacts caused by the time lag in database updates. In addition, annual coverages to PDB entries were calculated to determine fitting time ranges in each fold database.

For membrane proteins, we retrieved annual statistics from the mpstruc database, a curated resource of membrane proteins with known three-dimensional structures, as of April 2026.

* The latest version of the original PDB-only ECOD

### Comparison with other S-shaped curves

As described in the *Results* section, the Gompertz function was evaluated by comparison with the S-shaped functions such as the Logistic function (eq. 2), the Gaussian CDF (eq. 3), and the Weibull CDF (eqs. 4 and 5), defined as presented below.

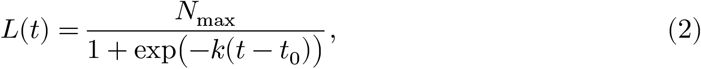

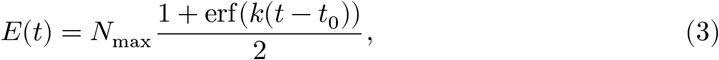

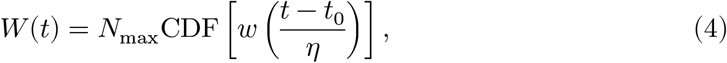

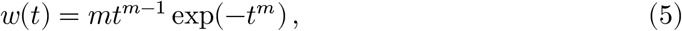

Therein, all parameters *N*_max_, *t*_0_, *k*, σ, and *m* serve as regression parameters with *N*_max_ representing the upper limit. Because the Weibull CDF is defined only for *t* > *t*_0_, parameter *t*_0_ was fixed as 1975, where the first structure was submitted the following year.

### Gompertz plot for the PDB growth curve

Taking the double logarithm on both sides of Gompertz definition (eq. 1), while accounting for the sign and the constant term, leads to a linear dependence on the time variable *t*:

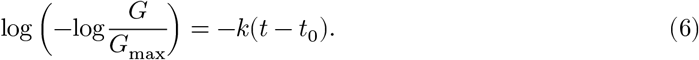

This linearity confirms that the growth curve follows the Gompertz function.

### Weibull plot for the PDB growth curve

To validate growth curve fitting using the Weibull CDF, we constructed a Weibull plot using the following transformations

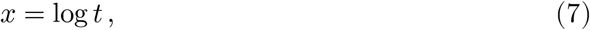

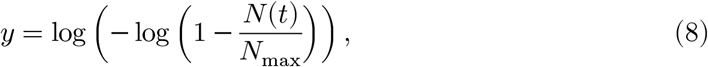

where *N* (*t*) denotes the growth curve of the protein structural database such as the total number of PDB entries, and where *N*_max_ represents the maximum of *N* (*t*). For the Weibull CDF (obtained with the time shift *t* − *t*_0_ → *t* in eqs. 4 and 5)

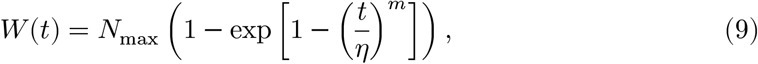

this transformation yields the linear form shown below:

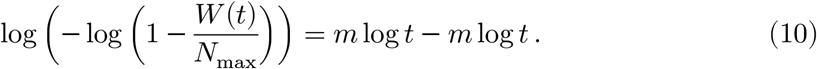

Therefore, the Weibull plot is expected to be a straight line with slope *m*, representing the order of the initial growth rate, and intercept −*m* log *η*.

### Fitting Methods

Curve fittings and evaluations were performed using Python, primarily with the SciPy library.

## Data Availability

All datasets are available at Zenodo repository (https://doi.org/10.5281/zenodo.19381698).

## Acknowledgements

This research was supported by Research Support Project for Life Science and Drug Discovery (Basis for Supporting Innovative Drug Discovery and Life Science Research (BINDS)) from AMED under Grant Number JP26ama121028.

## Author contributions

K. T. conceived the project. K. S. & K. T. contributed to the design and interpretation of this study. K. S. performed statistical analysis. K. S. & K. T. wrote the manuscript.

