## Supplementary Information for "The Gompertz curve for estimating growth rates of Protein Data Bank and protein folds"

### Contents

**Supplementary Note S1.** Logit plot for the growth curve
**Supplementary Note S2.** Probit plot for the growth curve
**Supplementary Note S3.** Convergence of Extreme Rarity to the Gumbel Law
**Supplementary Table S1.** Parameters of the Logistic function
**Supplementary Table S2.** Parameters of the Gaussian CDF
**Supplementary Table S3.** Parameters of the Weibull CDF
**Supplementary Table S4.** Fitting results for X-ray archive
**Supplementary Table S5.** Fitting results for NMR archive
**Supplementary Table S6.** Fitting results for EM archive
**Supplementary Figure S1.** Growth curve of the PDB archive (expanded view of Fig. 1B)
**Supplementary Figure S2.** Growth curve of mpstruc archive
**Supplementary References**

**Supplementary Note S1.** Logit plot for the growth curve.

The logit transformation, defined for $0 \leq p \leq 1$ as

$$\mathrm{logit}(p) = \log \frac{p}{1-p}, \tag{S1}$$

is the inverse function of the logistic function. A logit plot with time on the horizontal axis and the logit values on the vertical axis becomes linear if the growth follows a logistic form. For the growth curves analyzed in the main text, the same transformation is applied after normalization by the scale factor.

**Supplementary Note S2.** Probit plot for the growth curve.

The probit transformation, defined for $0 \leq p \leq 1$ as

$$\mathrm{probit}(p) = \Phi^{-1}(p), \tag{S2}$$

where $\Phi$ is the standard Gaussian cumulative distribution function (Gaussian CDF), is its inverse. Accordingly, a plot with time on the horizontal axis and the probit values on the vertical axis becomes linear if the growth follows a Gaussian cumulative distribution.

**Supplementary Note S3.** Convergence of Extreme Rarity to the Gumbel Law

#### Extreme Value Convergence to Gumbel

Consider a model in which each element is assigned to a category (such as protein fold groups) according to an underlying probability distribution, and categories are discovered through the exploration of the elements.

Let $\lambda$ denote the assignment probability of a category, and define the rarity

$$X = -\log \lambda.$$

Assume that $\{X_i\}_{i=1}^{n}$ are independent and identically distributed random variables, consider the maximum

$$X_n^{\max} = \max_{1 \le i \le n} X_i.$$

If the tail probability $\Pr(X > x)$ decays exponentially or faster as $x \to \infty$, then $X$ belongs to the Gumbel domain of attraction. Consequently, by the Fisher–Tippett–Gnedenko theorem, the suitably normalized maximum $X_n^{\max}$ converged in distribution to the Gumbel distribution.

#### Application to Protein Fold Rarity

The probability density function of the protein fold distribution assumed by Govindarajan et al. is given by

$$\rho(\lambda) = C \exp(-\alpha \lambda^{\beta}),$$

where $C$ is a normalization constant and $\alpha$ and $\beta$ are parameters.[1]

As the exploration of folds progresses, the contribution from rare fold ($\lambda \ll 1$) becomes dominant. In this regime, the density approaches a constant value, $\rho(\lambda) \sim C$ and the corresponding tail behavior of the rarity ($X = -\log \lambda$) is given by

$$\Pr(X > x) = \int_0^{e^{-x}} \rho(\lambda)\, \mathrm{d}\lambda \sim C \exp(-x),$$

showing an exponential tail.

In this case, choosing the normalization constants

$$a_n := 1, \quad b_n := \log(Cn),$$

one obtains

$$\begin{aligned}
\Pr\left(\frac{X_n^{\max} - b_n}{a_n} \le z\right) &= [F(a_n z + b_n)]^n \\
&= \left[1 - C\exp\bigl(-(z + \log(Cn))\bigr) + o\left(\frac{1}{n}\right)\right]^n \quad (n \to \infty) \\
&\sim \left[1 - \frac{\exp(-z)}{n}\right]^n \\
&\to \exp(-\exp(-z))\,,
\end{aligned}$$

which shows that the suitably normalized maximum converges in distribution to the Gumbel distribution.

In contrast, convergence to a Weibull distribution arises when the rarity $X$ has a finite upper bound. That is, there exists an endpoint $x_F$ such that

$$X \le x_F \le \infty,$$

and the tail behavior near the endpoint satisfies

$$\Pr(X > x_F - \epsilon) \sim C\epsilon^{\alpha} \quad (\epsilon \searrow 0).$$

However, current observations do not suggest the existence of a clear upper limit to the difficulty of protein fold discovery. In particular, a scenario in which highly difficult fold groups are concentrated near a finite upper bound appears implausible. Such a picture is also inconsistent with previous studies, including those by Govindarajan et al.[1].

**Supplementary Table S1.** Parameters of the Logistic function.

| | $N_{\mathrm{max}}$ | $k$ / yr$^{-1}$ | $t_0$ / yr |
|---|---|---|---|
| PDB | $(3.48 \pm 0.14) \times 10^5$ | $(1.51 \pm 0.04) \times 10^{-1}$ | $(2.0198 \pm 0.0006) \times 10^3$ |
| X-ray | $(2.48 \pm 0.04) \times 10^5$ | $(1.69 \pm 0.02) \times 10^{-1}$ | $(2.0171 \pm 0.0003) \times 10^3$ |
| NMR | $(1.45 \pm 0.02) \times 10^4$ | $(2.07 \pm 0.07) \times 10^{-1}$ | $(2.0083 \pm 0.0002) \times 10^3$ |
| EM | $(7.0 \pm 0.4) \times 10^4$ | $(4.09 \pm 0.08) \times 10^{-1}$ | $(2.0255 \pm 0.0002) \times 10^3$ |

**Supplementary Table S2.** Parameters of the Gaussian CDF.

| | $N_{\mathrm{max}}$ | $\sigma$ / yr | $t_0$ / yr |
|---|---|---|---|
| PDB | $(4.29 \pm 0.19) \times 10^5$ | $(1.80 \pm 0.05) \times 10$ | $(2.0229 \pm 0.0007) \times 10^3$ |
| X-ray | $(2.71 \pm 0.03) \times 10^5$ | $(1.51 \pm 0.01) \times 10$ | $(2.0183 \pm 0.0001) \times 10^3$ |
| NMR | $(1.44 \pm 0.02) \times 10^4$ | $(1.14 \pm 0.03) \times 10$ | $(2.0082 \pm 0.0002) \times 10^3$ |
| EM | $(1.7 \pm 0.3) \times 10^5$ | $8.3 \pm 0.4$ | $(2.030 \pm 0.001) \times 10^3$ |

**Supplementary Table S3.** Parameters of the Weibull CDF.

| | $N_{\mathrm{max}}$ | $\eta$ / yr | $m$ | $t_0$ |
|---|---|---|---|---|
| PDB | $(4.19 \pm 0.24) \times 10^5$ | $(5.16 \pm 0.10) \times 10$ | $4.51 \pm 0.07$ | 1975 |
| X-ray | $(2.46 \pm 0.03) \times 10^5$ | $(4.53 \pm 0.02) \times 10$ | $5.02 \pm 0.04$ | 1975 |
| NMR | $(1.48 \pm 0.02) \times 10^4$ | $(2.35 \pm 0.03) \times 10$ | $2.68 \pm 0.06$ | 1988 |
| EM | $(7.0 \pm 0.8) \times 10^4$ | $(3.16 \pm 0.05) \times 10$ | $9.73 \pm 0.22$ | 1995 |

**Supplementary Table S4.** Fitting results for X-ray archive.

| | RMSE | AIC | Estimated Limit |
|---|---|---|---|
| Gompertz | $0.36 \times 10^3$ | $1.19 \times 10^3$ | $(4.34 \pm 0.04) \times 10^5$ |
| Logistic | $1.78 \times 10^3$ | $1.50 \times 10^3$ | $(2.48 \pm 0.04) \times 10^5$ |
| Gaussian CDF | $0.63 \times 10^3$ | $1.29 \times 10^3$ | $(2.71 \pm 0.03) \times 10^5$ |
| Weibull CDF | $0.75 \times 10^3$ | $1.33 \times 10^3$ | $(2.46 \pm 0.03) \times 10^5$ |

**Supplementary Table S5.** Fitting results for NMR archive.

| | RMSE | AIC | Estimated Limit |
|---|---|---|---|
| Gompertz | $1.51 \times 10^2$ | $7.49 \times 10^2$ | $(1.63 \pm 0.02) \times 10^4$ |
| Logistic | $3.28 \times 10^3$ | $8.63 \times 10^3$ | $(1.45 \pm 0.02) \times 10^4$ |
| Gaussian CDF | $2.69 \times 10^2$ | $8.34 \times 10^2$ | $(1.44 \pm 0.02) \times 10^4$ |
| Weibull CDF | $2.05 \times 10^2$ | $7.94 \times 10^2$ | $(1.48 \pm 0.02) \times 10^4$ |

**Supplementary Table S6.** Fitting results for EM archive.

| | RMSE | AIC | Estimated Limit |
|---|---|---|---|
| Gompertz | $1.91 \times 10^2$ | $6.36 \times 10^2$ | $(9.9^{+4.8}_{-3.3}) \times 10^5$ |
| Logistic | $1.49 \times 10^2$ | $6.07 \times 10^2$ | $(7.0 \pm 0.4) \times 10^4$ |
| Gaussian CDF | $1.76 \times 10^2$ | $6.27 \times 10^2$ | $(1.7 \pm 0.3) \times 10^5$ |
| Weibull CDF | $1.72 \times 10^2$ | $6.24 \times 10^2$ | $(7.0 \pm 0.8) \times 10^4$ |

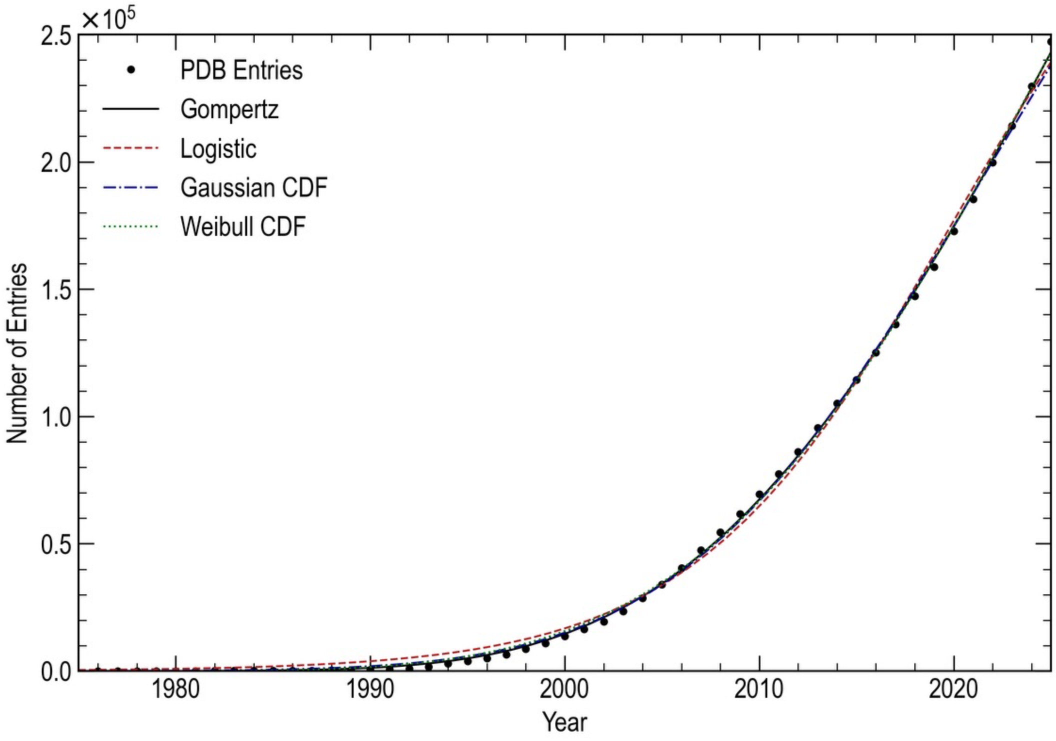


**Supplementary Figure S1.** Expanded view of Fig. 1B showing the growth curve of the PDB archive with comparison of the Gompertz fit (black) to other sigmoidal functions, including the logistic function (red), the Gaussian cumulative distribution function (Gaussian CDF) (blue), and the Weibull cumulative distribution function (Weibull CDF) (green).

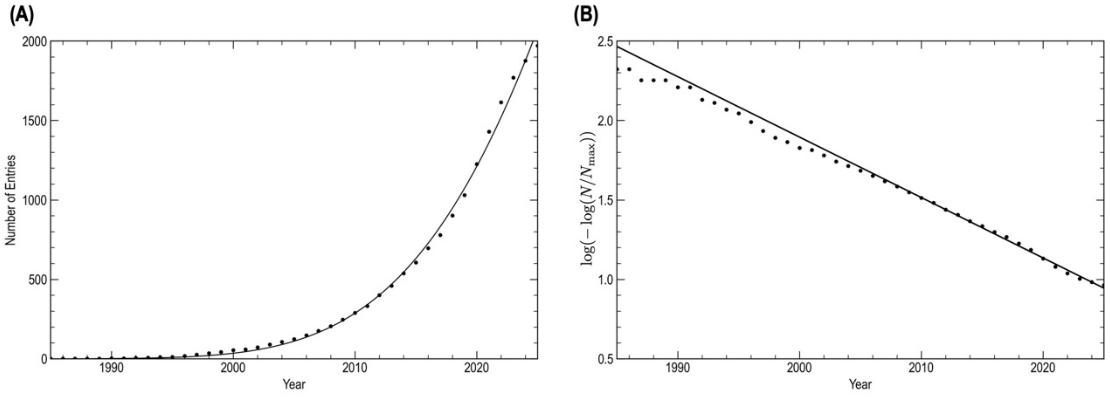


**Supplementary Figure S2.** Growth curve of mpstruc archive. Statistical data were obtained from mpstruc[2]. (A) Growth curve of the total mpstruc entries (dots) fitted with the Gompertz function (line). (B) Gompertz plot for the mpstruc growth.
